# Detecting Early Response to Immune Checkpoint Blockade by Multimodal Molecular Imaging

**DOI:** 10.1101/2020.04.30.070508

**Authors:** Yu Saida, Jeffery R. Brender, Kazutoshi Yamamoto, James B. Mitchell, Murali C. Krishna, Shun Kishimoto

## Abstract

Immune checkpoint inhibitors have become a standard therapy for several cancers; however, the response is inconsistent and a method for non-invasive assessment has not been established to date. To investigate the capability of multi-modal imaging to evaluate treatment response to immune checkpoint blockade therapy, we employed hyperpolarized ^13^C MRI on tumor bearing mice using [1-^13^C] pyruvate and [1,4-^13^C_2_] fumarate to detect early changes in tumor glycolysis and necrosis, respectively. Following αPD-L1 Ab + αCTLA-4 Ab dual immune checkpoint blockade (ICB) therapy, dynamic contrast enhanced (DCE) MRI was used to determine the treatment effect on intratumor perfusion/permeability. Mice bearing MC38 colon adenocarcinoma and B16.F10 melanoma were used as sensitive and less sensitive models, respectively to immune checkpoint dual blockade of PD-L1 and CTLA-4. Glycolytic flux significantly decreased upon treatment in the less ICB sensitive B16.F10 model but remained essentially unchanged in MC38 tumors. Imaging [1,4-^13^C] fumarate conversion to [1,4-^13^C] malate showed a significant increase in necrosis in the treatment group for the ICB sensitive MC38 tumor (*p* = 0.0003), with essentially no change in ICB sensitive B16.F10 tumors. Histological assessment showed increased necrotic tissue with enhanced lymphocyte infiltration in the MC38 treatment group, suggesting immunogenic tumor cell death. Dynamic contrast enhanced MRI showed significantly increased perfusion/permeability of Gd-DTPA in MC38 treated tumor, while a similar trend but statistically non-significant change was observed in B16.F10 treated tumor. These results provide imaging biomarkers to detect early response to cancer immunotherapy, allowing qualitative assessment of tumors treated with immune checkpoint blockade therapy.

## Introduction

Immunotherapy using immune checkpoint blockade (ICB) has emerged as a promising cancer treatment^1^. ICB therapy removes inhibitory signals of T cell activation by blocking co-stimulatory receptors or ligands such as PD-1, PD-L1, and CTLA-4, enabling tumor-reactive T cells to override regulatory mechanisms and exhibit antitumor activity^2^. A key feature of this therapy is the highly durable tumor response, resulting in a plateau in the tail of the survival curve unlike chemotherapy or genomically targeted therapy the benefits of which tend to diminish as time progresses^3^. Despite the potential of this type of therapy, the remarkable responses to ICB are limited to a fraction of patients. Thus, an accurate and reliable tool for assessment of early treatment response is needed. While several biomarkers including PD-L1 expression, tumor mutation burden, mismatch repair deficiency, tumor-infiltrating lymphocytes (TILs) have been proposed, none of them has become a gold standard for predictor of treatment response to date^4^. The reason for this absence is explained by the unique response to immunotherapy. Unlike cytotoxic therapy, where the effectiveness can be frequently judged by the reduction in the size of the tumor, the treatment response in immunotherapy is often delayed; tumors transiently enlarge or new lesions appear in the early stages only to be followed by tumor shrinkage or long-term stability of tumor size^5,6^. The delayed response can pose a potential problem in designing treatment plans. In order to alleviate this problem, the immune-related response criteria (irRC), immune-related response criteria (irRECIST), and immune RECIST (iRECIST) were developed for imaging assessment of treatment response. These criteria require a consecutive scan at least 4 weeks apart for confirmation of progressive disease^6,7^. When the treatment is found ineffective, patients may lose the opportunity to receive other treatments during the assessing time. Thus, developing an early prediction of the treatment response to immunotherapy is of great importance^8^.

Treatment response to cancer immunotherapy is closely related to the tumor microenvironment. Immunologically “hot” tumors are typically characterized by a high degree of CD8+ cytotoxic T cells infiltration^9^. A lack of CD8+ T cell infiltration and the presence of immune suppressor cells including regulatory T cells, myeloid derived suppressor cells, and type II macrophages are one of extrinsic mechanisms of resistance to immunotherapy^10^. Recently, increased metabolic competition between tumor cells and immune cells was reported. A metabolic effect characteristic of cancer cells, aerobic glycolysis, also called “Warburg effect” (Warburg 1956) causes depletion of extracellular glucose and restricts glucose availability to T cells leading to suppression of glycolytic metabolism in T cells, resulting in decreased effector function^11^. In this context, it is reported that PD-L1 promotes glycolysis in tumors via Akt/mTOR activation, and that PD-1 blockade can decrease glycolysis in tumors^12^. It is also well known that tumor cells create a challenging environment for the immune system characterized by hypoxia, low pH, high interstitial fluid pressure, and immune-inhibitory metabolites such as lactic acid^13,14^. The impaired perfusion capacity of tumor blood vessels by abnormal tumor vasculature create a highly hypoxic microenvironment, frequently resulting in immune suppression^15^. Recent studies have demonstrated that tumor hypoxia is negatively associated with the efficacy of immunotherapy and that the stimulation of immune cell functions with PD-1 and CTLA-4 blockade can also help to normalize tumor vessels^16,17^. In this regard, it is potentially feasible to assess the treatment efficacy of immunotherapy by evaluating the metabolism and microenvironment of the tumors before a detectable reduction in the size of the tumor.

A few studies have reported that ^18^F-FDG-PET/CT can potentially predict early response to PD-1 blockade or CTLA-4 blockade in patients with non-small cell lung cancer or melanoma^18,19^. However, several studies have also demonstrated the inability of ^18^F-FDG-PET/CT to distinguish patients with pseudoprogression from those with progressive disease in melanoma patients^20,21^. Thus, to date, non-invasive imaging approaches to clarify the early response to effective immunotherapy in vivo have not been established.

The purpose of this study is to investigate the capability of non-invasive metabolic and physiologic imaging to evaluate early response to immune checkpoint blockade therapy. New advancements in imaging using dynamic nuclear hyperpolarization (DNP) is used to enhance ^13^C signal intensity in vivo by approximately four orders of magnitude. By using ^13^C pyruvate as a probe, it is possible to monitor the glycolytic profile of a tumor non-invasively, which has proven useful for monitoring the treatment response to several chemotherapeutics such as anti- angiogenic agent sunitinib and hypoxia-activated prodrug TH-302^22,23^. Similarly, the [1,4-^13^C_2_] fumarate probe is expected to detect necrotic tumor cell death in vivo^24,25^.

To test the applicability of these new imaging techniques along with conventional dynamic contrast enhanced (DCE) MRI in detecting ICB treatment response, we imaged mice bearing murine colon adenocarcinoma carcinoma (MC38) tumor or melanoma (B16.F10) tumor as ICB sensitive and less ICB sensitive models, respectively. Using the combination therapy of PD-L1/PD-1 and CTLA-4 as a treatment model, which has improved therapeutic effect compared with either monotherapy^26–28^, multimodal metabolic and physiologic imaging showed substantial differences between the sensitive and less sensitive models, which may serve as a guide for further development of imaging biomarkers.

## Results

### In vivo treatment effect of immune checkpoint inhibitors in murine tumor model

Anti-PD-L1 monocronal antibody (mAb) monotherapy or in combination with anti-CTLA-4 mAb was used to evaluate the therapeutic effect immune checkpoint blockade in vivo. Mice were inoculated subcutaneously with either MC38 or B16.F10 cells (1 × 10^5^) into the right leg and treated with either or both immune checkpoint inhibitors by i.p. injection every three days for a total of three injections.

By itself, anti-PD-L1 Ab therapy inhibited tumor growth in MC38 colon adenocarcinoma tumors (Fig. 1a) and extended survival times (Fig. 1b, 1c) compared to the control group (median survival time 29.5 days compared to 24 days without treatment, *p* = 0.0003). Combining anti-PD-L1 with anti-CTLA-4 Ab treatment resulted in stronger inhibition of tumor growth and a statistically significant increase in survival time relative to anti-PD-L1 Ab alone (33.5 days to 29.5 days, *p* = 0.0475, Fig. 1b). Both anti-PD-L1 Ab by itself and the combination treatment anti-PD-L1 Ab + anti-CTLA-4 Ab also delayed tumor growth (Fig. 1d) and extended survival in B16.F10 tumors (median survival time 22 days for anti-PD-L1 Ab and 24 for the combination treatment compared to 19.5 days without treatment*, p* = 0.0071, *p* = 0.0018, respectively, Fig. 1e, 1f). However, the cytoreductive effect of anti-PD-L1 Ab itself was less in B16.F10 than in MC38 tumors and the combination treatment with anti-CTLA-4 Ab did not significantly prolong survival time compared to anti-PD-L1 Ab alone (24 days compared to 22 days, *p* = 0.0752, Fig. 1e). These results are consistent with previous reports showing that MC38 is sensitive to ICB therapy, while B16.F10 is less sensitive ^29^.

**Figure 1.**
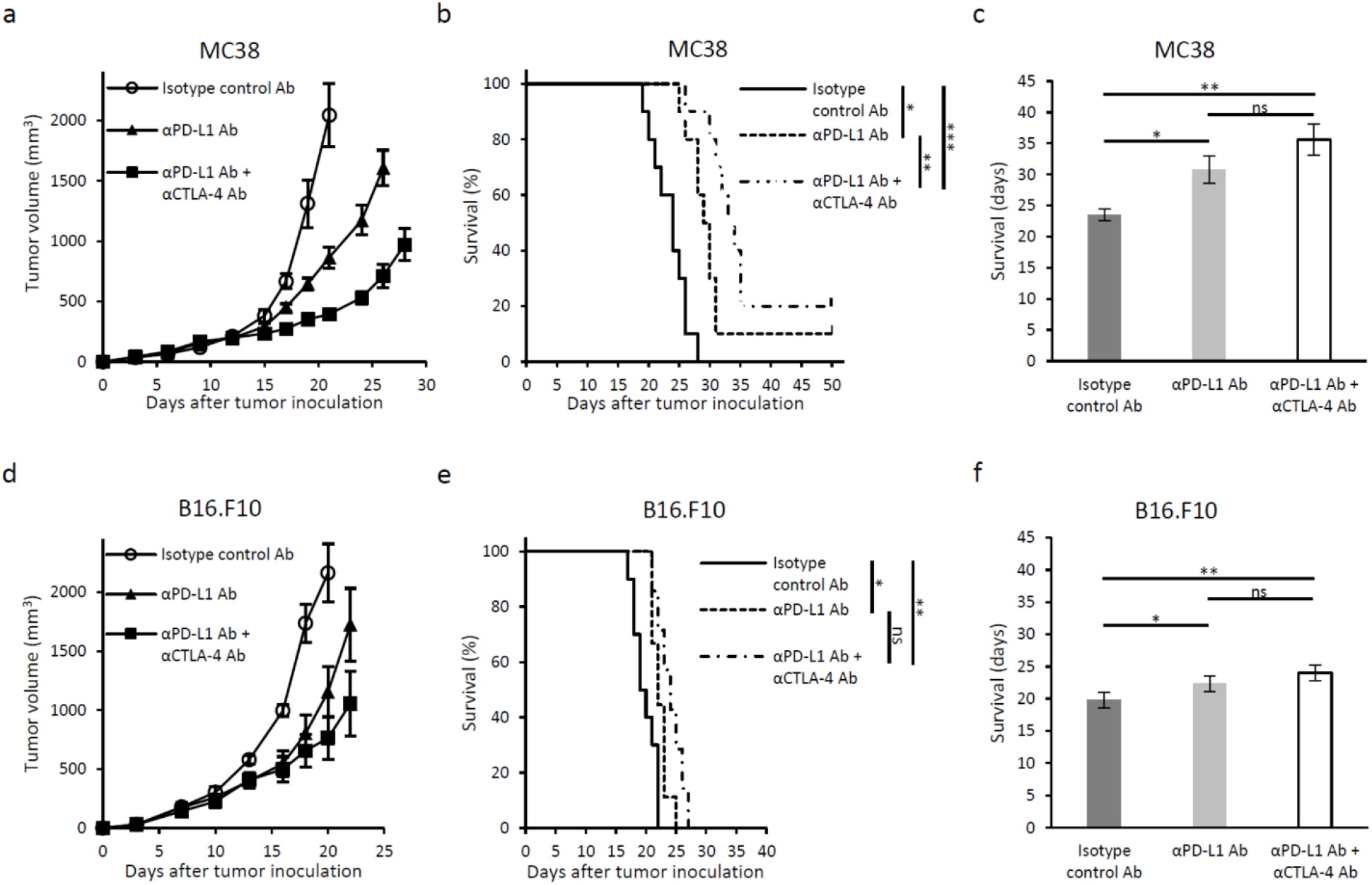
Tumor growth and survival of immune checkpoint blockade therapy in two murine tumor models with different sensitivity. **a** and **d,** Growth kinetics of each tumor. MC38 or B16.F10 inoculated mice were treated with either Isotype control Ab, anti-PD-L1 Ab, or combination anti-PD-L1 Ab + anti-CTLA-4 Ab on days 9, 12, and 15 post tumor inoculation (n = 5 per group). Data are shown as mean ± SE at each time point. **b** and **e,** Kaplan-Meier survival curve for each tumor (MC38; n = 10 per group, B16.F10; n =7-10 per group). Survival refers to the time before reaching the maximally allowed tumor volume of 2,000 mm^3^. The differences between groups in the Kaplan-Meier curve are identified using the log-rank test (**b,** \**p* = 0.0003, \*\**p* = 0.0475, \*\*\**p* < 0.0001 **d,** \**p* = 0.0071, \*\**p* = 0.0018). **c** and **f,** Bar plot of survival from figure B and E. Data are shown as mean ± SE (**c,** \**p* = 0.0427, \*\**p* = 0.0007, **f,** \**p* = 0.0140, \*\**p* = 0.0002).

This difference in sensitivity to ICB therapy was not reflected in T-cell counts (Fig. 2a). Intratumor levels of CD3+CD8+ T cells increased substantially in both tumor models after 4 days of anti-PD-L1 Ab + anti-CTLA-4 Ab combination treatment, suggesting effective infiltration of lymphocytes into the tumors. Combination treatment only raised levels of CD3+CD4+ and CD3+CD8+ T cells in tumor draining lymph nodes in MC38 tumors, but the populations of both types of cells was already at a similar level in the absence of treatment in B16.F10. To more quantitatively measure total and activated T cells in situ, histological assessment was performed on MC38 tumor slices to confirm T cells proliferation. Immunofluorescence staining of CD3 and CD8 showed that infiltration of CD3+ T cells and CD8+ cytotoxic T cells significantly increased in anti-PD-L1 Ab and anti-CTLA-4 Ab treated tumor compared to isotype control Ab treated tumor (*p* < 0.0001, *p* < 0.0001), confirming an active immune response as early as day 4 after the combination ICB treatment in both tumor models (Fig. 2b, 2c). The CD8 to CD3 ratio was also similar between groups suggesting a similar level of activation between CD4 and CD8 T cells (Fig. 2c). Although T cell activation is a complex process subject to multiple points of regulation, the data suggest a simple measure of CD8+ and CD4+ T cell counts is not sufficient to predict ICB efficacy.

**Figure 2.**
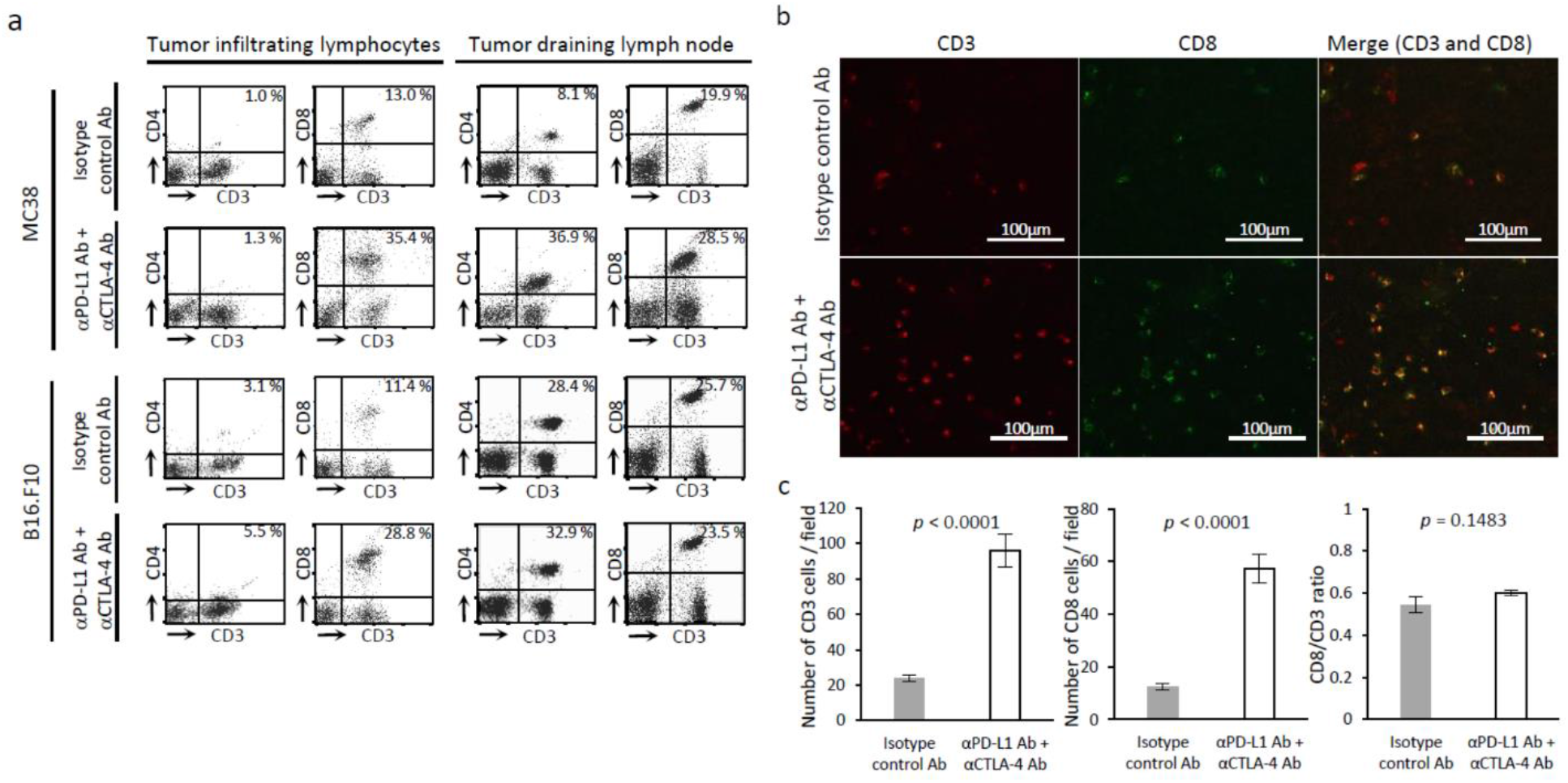
In vivo treatment effect of immune checkpoint blockade therapy. **a,** Flow cytometry analysis of tumor infiltrating lymphocytes and tumor draining lymph node cells. Tumor bearing mice were treated with each Ab on day 9 and 12. Tumors and lymph nodes were harvested on day 13. The figure on each plot shows the percentage of CD3+CD4+ T cells or CD3+CD8+ T cells. **b** and **c,** Immunofluorescence staining of CD3 and CD8 in MC38 tumor (scale bar = 100 μm). MC38 tumor bearing mice were treated with each Ab on day 9 and 12. Tumors were harvested on day 13. Number of cells were manually counted in randomly selected 5 fields per tumor section (n = 4 per group). Data are shown as mean ± SE.

### Changes in tumor glycolytic metabolism induced by immune checkpoint blockade

Activation, proliferation, and differentiation of T cells all require the synthesis of numerous macromolecules. To generate synthetic intermediates to support these processes, naïve T cells often undergo metabolic reprogramming to enhance glycolytic flux. Decreases in glycolytic flux therefore could serve as a potential biomarker to predict ICB efficacy. To investigate whether glycolytic metabolism of tumor cells is altered directly by PD-L1 blockade, we measured the *in vitro* extracellular acidification rate (ECAR) of tumor cells by the Seahorse assay to determine the extent of lactic acid fermentation (Fig. 3a). The ECAR of MC38 cells in vitro was not altered after treatment with anti-PD-L1 Ab, except for a slight and statistically insignificant increase in baseline acidification not attributable to glycolysis. In contrast, the ECAR of the less ICB sensitive B16.F10 cell line was significantly reduced after treatment with anti-PD-L1 Ab (Fig. 3b), suggesting that the glycolytic metabolism of B16.F10 is more dependent on the PD-L1/PD-1 pathway than that of MC38. To investigate whether expression of glycolytic enzymes and transporters are affected by PD-L1 blockade, PD-L1 inhibitor treated tumor cells were analyzed by western blotting and flow cytometry. No significant changes in the expression of glycolytic enzymes and transporters after treatment were detected in either cell line (Supplementary Fig. 1).

**Figure 3.**
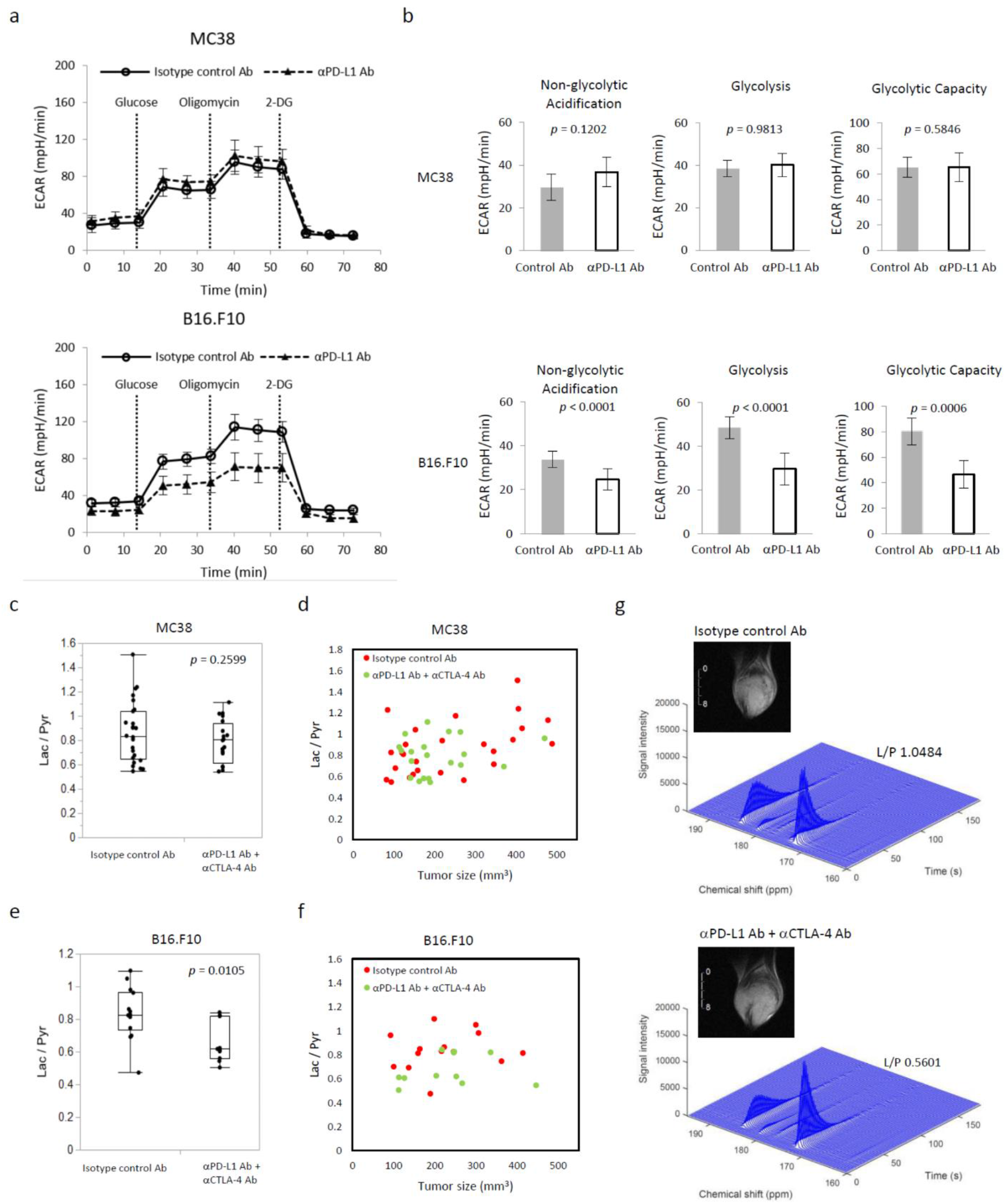
Metabolic shift induced by immune checkpoint blockade can be detected in selected cancer whose metabolism relies on glycolysis. **a,** Extracellular acidification rate (ECAR) of MC38 and B16.F10 cells in vitro. Tumor cells pre-treated with IFN-◻ for 48 h followed by anti-PD-L1 Ab or isotype control Ab for 24 h were measured (n = 5 per group). For each time point, mean ± SD is plotted. **b,** Parameters (non-glycolytic acidification, glycolysis, and glycolytic capacity) calculated from ECAR kinetics. Data are shown as mean ± SD. **c-g,** MRI of hyperpolarized ^13^C pyruvate metabolism in two murine tumor models. Tumor bearing mice treated with isotype control Ab or anti-PD-L1 Ab + anti-CTLA-4 Ab were scanned after treatment (MC38; n = 26, n = 20 each group, B16.F10; n = 14, n = 11 each group). **c,** Lactate to Pyruvate (Lac/Pyr) ratio of MC38 sorted by treatment. Data are shown as box-and-whisker plot (median, maximum, minimum, first quartile, and third quartile); individual values are shown. **d,** Correlation between Lac/Pyr ratio and tumor size. **e,** Lac/Pyr ratio of B16.F10 sorted by treatment. Data are shown as box-and-whisker plot and individual values. **f,** Correlation between Lac/Pyr ratio of B16.F10 and tumor size. **g,** Representative dynamic ^13^C spectra of B16.F10 tumor and T2-weighted ^1^H anatomical image.

To better understand the metabolic changes associated with ICB, we measured the flux through the lactate dehydrogenase pathway in vivo by hyperpolarized ^13^C MRI using [1-^13^C] pyruvate. Non-localized 1D spectra encompassing the entire tumor implanted leg were acquired continuously for 240 s after the injection of hyperpolarized [1-^13^C] pyruvate and the conversion of pyruvate to lactate was quantitatively evaluated (Fig. 3g). The lactate/pyruvate ratio is a direct measure of flux through the lactate dehydrogenase pathway and an indirect measure of glycolysis. No difference in the median Lac/Pyr ratio was detected in the ICB sensitive MC38 tumors between the anti-PD-L1 Ab + anti-CTLA-4 Ab treatment and isotype control Ab groups by the final time point (median Lac/Pyr ratio 0.8039 for the combination treatment compared to 0.8312 for the control, *p* = 0.2599) (Fig. 3c), although a cluster of large tumors with high Lac/Pyr ratios likely undergoing anaerobic fermentation is evident in the control group (Fig. 3d). No correlation of the Lac/Pyr ratio with the treatment group was found when controlled for tumor size to remove the effect (*p* = 0.3737, Supplementary Table 1a). Results from the ICB insensitive B16.F10 tumors are conspicuously different. The Lac/Pyr ratio was significantly lower in the in anti-PD-L1 Ab + anti-CTLA-4 Ab treatment group relative to the control group (median Lac/Pyr ratio 0.6178 for the combination treatment compared to 0.826 for the control, *p* = 0.0105, Fig.3e). Even when controlled for tumor size, the Lac/Pyr ratio remained negatively correlated with the treatment with statistical significance (*p* = 0.0246, Supplementary Table 1b and Fig. 3f). Taken together, these results suggest the relationship between PD-L1 blockade and glycolysis is not straightforward and is subject to the control of the tumor microenvironment.

### Necrotic change induced by immune checkpoint blockade

Tumor cell necrosis is potentially a more direct biomarker for ICB efficacy. To evaluate necrotic cell death in tumors, hyperpolarized ^13^C MRI using [1,4-^13^C_2_] fumarate was performed on the same sets of tumors. Fumarate is not actively transported into the cell and the plasma membrane is impermeable to fumarate when intact. A loss of membrane integrity during necrosis allows entry of fumarate and the conversion to malate by cytosolic fumarate hydratase, which can be detected as a doublet distinct from fumarate at 182-183 ppm (Fig. 4c). Fig. 4a shows the malate to fumarate (Mal/Fum) ratio of ICB sensitive MC38 tumor bearing mice sorted by treatment group. Mal/Fum ratios were significantly higher in anti-PD-L1 Ab + anti-CTLA-4 Ab treated MC38 tumors relative to the control gtoup (median Fum/Mal ratio 0.0796 for the combination treatment compared to 0.058 for the control, *p* = 0.0003, Fig. 4a). The Mal/Fum ratio remained a significant variable even when tumor size was controlled for (*p* = 0.0037, Supplementary Table 2a and Fig. 4b), suggesting that tumor cell necrosis is induced as a treatment effect of ICB therapy *in vivo*. No statistically significant difference was found in the ICB insensitive B16.F10 tumors (median Fum/Mal ratio 0.0927 for the combination treatment compared to 0.0872 for the control, *p* = 0.4162, Fig. 4d). Multiple regression analysis confirmed that Mal/Fum ratio was not a significant predictor of the treatment group (*p* = 0.4465, Supplementary Table 2b). No difference in serum fumarase activity was found between ICB treated MC38 mice and the control (*p* = 0.7233, Fig.4f), suggesting that serum fumarase activity cannot be a surrogate marker to monitor ICB efficacy.

**Figure 4.**
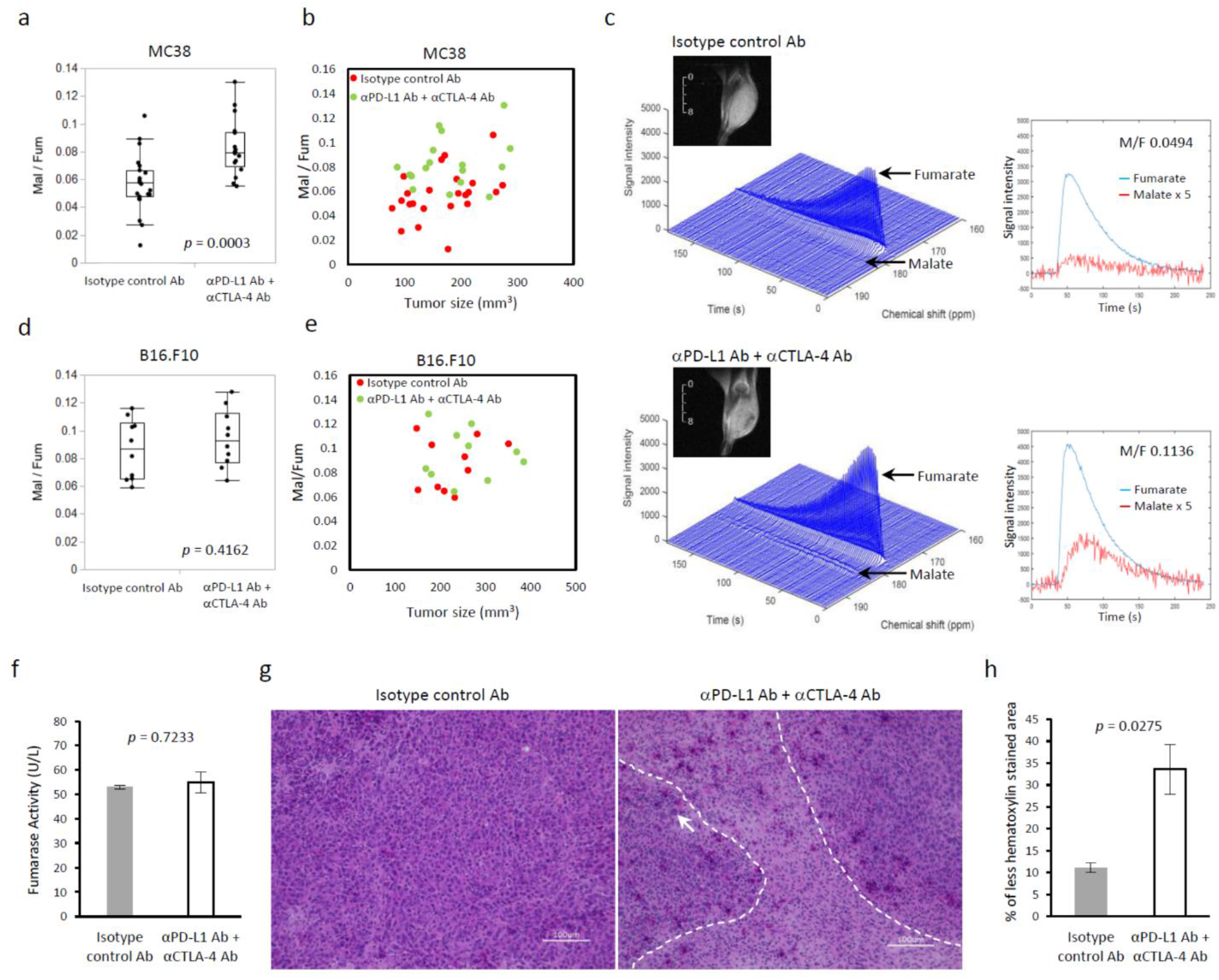
Tumor cell death can be detected by ^13^C fumarate MRI as treatment effect of immune checkpoint blockade. **a-e,** MRI of hyperpolarized ^13^C fumarate metabolism in two murine tumor models. Tumor bearing mice treated with isotype control Ab or anti-PD-L1 Ab + anti-CTLA-4 Ab were scanned after treatment (MC38; n = 23, n = 17 each group, B16.F10; n = 10 per group). **a,** The Malate to Fumarate (Mal/Fum) ratio of MC38 sorted by treatment. Data are shown as box-and-whisker plot (median, maximum, minimum, first quartile, and third quartile); individual values are shown. **b,** Correlation between Mal/Fum ratio and tumor size. **c,** Representative dynamic ^13^C spectra of MC38 tumor and T2-weighted ^1^H anatomical image. Signal of Malate is quintupled in the right plot. **d,** Mal/Fum ratio of B16.F10 sorted by treatment. Data are shown as box-and-whisker plot and individual values. **e,** Correlation between Mal/Fum ratio of B16.F10 and tumor size. **f,** Fumarase activity assay of mice serum. MC38 tumor bearing mice were treated with isotype control Ab or anti-PD-L1 Ab + anti-CTLA-4 Ab on day 9 and 12. Blood were collected on day 13. Data are shown as mean ± SE (n = 5, n = 7 each group). **g,** Representative hematoxylin and eosin staining (HE) of MC38 tumor (scale bar = 100 μm). MC38 tumor bearing mice were treated with each Ab on day 9 and 12. Tumors were harvested on day 13. **h,** Quantification of less hematoxylin stained area. Data are shown as mean ± SE (n = 4 per group).

To confirm the necrotic change by histology, tumor sections were stained with hematoxylin and eosin according to standard procedures. In the ICB sensitive MC38 tumors, infiltration of lymphocytes was confirmed by focal areas of hematoxylin negative spaces (dot-line area, Figure 4g). Nucleic enlargement in tumor cells (white arrow) indicating necrotic cell damage was also observed (Fig. 4g). The overall decrease in hematoxylin positive area, indicating the replacement with interstitial tissue after tumor cell death, was quantified using ImageJ software, and found to be significantly increased in anti-PD-L1 Ab + anti-CTLA-4 Ab treated tumor compared to the control (*p* = 0.0275, Fig. 4h). These histologic findings support the necrotic cell death as a consequence of antitumor effect of lymphocytes induced by ICB therapy.

### Changes in Intratumor blood perfusion induced by immune checkpoint inhibitors

Tumor perfusion/permeability changes in response to ICB treatments were investigated by DCE-MRI using the Toft model. In the Toft model, permeability changes are characterized by the influx forward volume transfer constant (K^trans^) from plasma into the extravascular-extracellular space (EES), which reflects the sum of all processes (predominantly blood flow and capillary leakage) that determine the rate of gadolinium influx from plasma into the EES. We examined isotype control Ab treated and anti-PD-L1 Ab + anti-CTLA-4 Ab treated tumor bearing mice after the 2^nd^ Ab injection. Relative to the control, Gd-DTPA uptake was increased (Fig. 5a and 5b) and K^trans^ was significantly higher in anti-PD-L1 Ab + anti-CTLA-4 Ab treated MC38 tumors (*p* < 0.0001, Fig. 5c). The separate effects of perfusion and permeability can be distinguished by the AUC of the calculated GD concentration at short (1 min) and long (10 min), respectively. Significant increases to both perfusion and permeability were observed (*p* < 0.0001, *p* < 0.0001, Fig. 5d). Changes in the ICB less sensitive B16.F10 tumors were substantially less. Perfusion (AUC 1 min) and K^trans^ increased slightly, although the difference was not statistically significant in either case (*p* = 0.3607 and *p* = 0.3076, respectively Fig. 5e and 5f). No difference in permeability was detected (AUC 10 min, 5f).These findings suggest that ICB treatment improved blood flow in MC38 tumor, but not in B16.F10.

**Figure 5.**
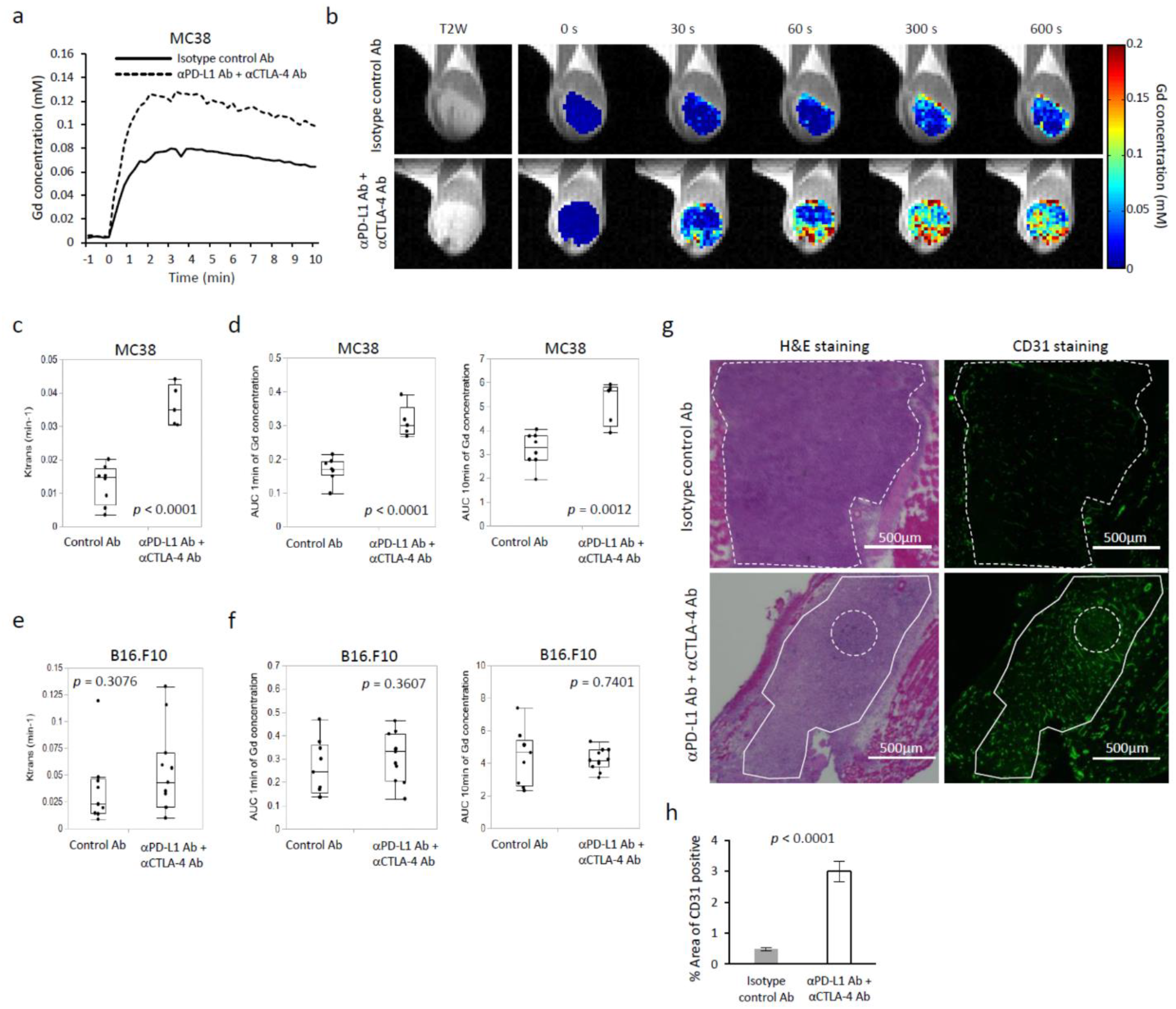
Tumor perfusion and permeability are improved by immune checkpoint blockade. **a-f,** Dynamic contrast enhanced MRI in two murine tumor model. Tumor bearing mice treated with isotype control Ab or anti-PD-L1 Ab + anti-CTLA-4 Ab were scanned after treatment (MC38; n = 8, n = 5 each group, B16.F10; n = 9, n = 11 each group). **a,** Representative time-intensity kinetic curve of Gd-DTPA in MC38 tumor. **b,** Gd-DTPA intensity with T2-weighted anatomical image of MC38 tumors. **c,** K^trans^ value of MC38 tumor sorted by treatment group. Data are shown as a box-and-whisker plot (median, maximum, minimum, first quartile, and third quartile); individual values are shown. **d,** Area under curve (AUC) 1 min and AUC 10 min of Gd-DTPA concentration in MC38. Data are shown as box-and-whisker plot and individual values. **e,** K^trans^ value of B16.F10 tumor sorted by treatment group. Data are shown as box-and-whisker plot and individual values. **f,** AUC 1 min and AUC 10 min of Gd-DTPA concentration in B16.F10. Data are shown as box-and-whisker plot and individual values. **g** and **h,** Immunofluorescence staining of CD31 in MC38 tumor. MC38 tumor bearing mice were treated with each Ab on day 9 and 12. Tumors were harvested on day 13 (n = 4 per group). **g,** Representative HE and CD31 staining of same section in MC38 tumor (scale bar = 500 μm). **h,** Quantification of CD31 positive area. Data are shown as mean ± SE (n = 4 per group).

To examine the changes in blood vessel density, IHC was performed with MC38 tumor sections. Fig. 5g shows the staining of CD31, which is a marker of vascular endothelial cells, as well as the staining of H&E in same section. Compared to Isotype control Ab treated tumor, CD31 staining was increased in anti-PD-L1 Ab + anti-CTLA-4 Ab treated tumor. CD31 positive area increased notably in hematoxylin less staining area where tumor cells are less (solid-line area), while CD31 positive area were poor in hematoxylin dense area where tumor cells are rich (dot-line area) in both control Ab and ICB treated tumor. Quantification of CD31 positive area showed significant increase in ICB treated tumor (*p* < 0.0001, Fig. 5h). These findings suggest improvement of vascular flow in MC38 tumor with ICB therapy, which supports the finding of increased tumor perfusion/permeability in DCE MRI.

## Discussion

In this study, we investigated the capability of non-invasive metabolic and physiologic imaging to evaluate the early response to ICB therapy in tumor bearing animal models. To our knowledge, this is the first report of metabolic imaging assessment to detect the early response to immune checkpoint blockade therapy using hyperpolarized ^13^C MRI.

Expression of PD-L1 on cancer cells promotes glucose uptake and production of lactate as part of the metabolic programming that enhances the survival of cells in hypoxic conditions and provides fuel for further growth. The exact mechanism by which this occurs is not fully understood. Previous reports have shown that blocking PD-L1 may directly dampen tumor glycolysis by inhibiting mTOR activity^13^. In MC38 and B16.F10 cell lines, PD-L1 blocking *in vitro* did not significantly alter expression of glycolytic enzymes and transporters (Supplementary Fig. 1). However, *in vivo* tumor metabolism when evaluated by ^13^C pyruvate MRI showed a significant decrease in glycolysis, but only in B16.F10 tumors where the effect of ICB on tumor volumes and survival times was substantially less. The reasons for this discrepancy are not entirely clear. The in vivo tumor microenvironment consists tumor cells with substantial contributions from the tumor stroma and immune cells. The expression of PD-1 on T cells is reported to inhibit glycolysis or amino acid metabolism and up-regulate fatty acid oxidation leading to impaired energy generation, which compromises proliferation and effector functions^30^. CTLA-4 can also cause decreased expression of GLUT1, increased mitochondrial oxidation and fatty acid uptake, and decreased biosynthesis on T cells^31,32^. Therefore, inhibition of PD-1/PD-L1 and CTLA-4 would preferentially promote T-cell function. In our study, histological assessments showed increased T cell population surrounding tumor cells in tissue. ICB treatment might restore the metabolic balance in favor of T cells, leading to a less glycolytic profile when the tumor is considered as a whole. In our study, glycolytic flux in MC38 tumor was not affected by ICB, but B16.F10 was significantly altered, in line with previous reports that show higher glycolytic activity in B16.F10 resulting in high acidification of the tumor microenvironment compared to MC38^33^. This suggested that B16.F10 is more dependent on glycolytic metabolism for energy production than MC38. Moreover, ECAR assays from previous reports^13^ and our data showed that anti-PD-L1 Ab treatment decreased ECAR more significantly in B16.F10 than in MC38, suggesting glycolytic metabolism in B16.F10 is more susceptible to anti-PD-L1 Ab. These findings suggested that the alteration of glycolytic metabolism was induced through interaction between tumor cells and tumor microenvironment including immune cells. In this respect, ^13^C pyruvate MRI may detect the effect of ICB therapy in selected cancers whose metabolism strongly relies on glycolysis.

^13^C fumarate MRI showed significantly increased Mal/Fum in ICB treated tumors relative to control tumors, suggesting necrotic tumor cell damage was induced in response to ICB therapy. The utility of the malate production from hyperpolarized [1,4-^13^C_2_] fumarate probe as a sensitive indicator of tumor cell death has been previously shown in several tumor types treated with a cytotoxic agent, anti-VEGF drug, or multi-kinase inhibitor^24,34,35^. In contrast to chemotherapy which directly shows cytotoxicity on tumor cells, the effect of ICB is induced via activation of immune system. Blockade of immune checkpoints such as PD-L1, PD-1, CTLA-4, whose signals help keep T cells from killing cancer cells, release the “brakes” on the immune system and enable T cells to activate and show cytotoxicity. There are two main mechanisms involved in cytotoxic T cells-mediated tumor cell death: one elicited by granule exocytosis (perforin and granzymes) and the other via the death ligand/death receptor system like Fas ligand (FasL) and TNF-related apoptosis inducing ligand (TRAIL). It has been assumed that the final consequence of granule exocytosis is the induction of cell death by apoptosis. However, recent experimental evidence indicating that perforin and granzymes of cytotoxic lymphocytes can activate non-apoptotic pathways of cell death^36^. Besides, recently, it has described that death receptor can induce “necroapoptosis”^37^. Necroapoptosis is a programmed form of necrosis, or inflammatory cell death, whereas necrosis is associated with unprogrammed cell death resulting from cellular damage. In necroapoptosis, rupture of the cell membrane and excretion of intracellular substrates occur as well as necrosis. Thus, we assume that mixed mechanisms of tumor cell death including a mechanism similar to necrosis allows ^13^C fumarate MRI to detect tumor cell death in response to ICB therapy in our study. Mal/Fum showed significant increase in the treated group of ICB sensitive MC38, and a much smaller change in the less ICB sensitive B16.F10 model. The results suggested that ^13^C fumarate MRI detects treatment effect of ICB therapy more accurately compared to ^13^C pyruvate MRI.

Treatment effects were also detectable by measurements of intratumor perfusion/permeability, by DCE MRI. Previous studies have shown that abnormal tumor vasculature contributes to immune suppression through multiple mechanisms. One such mechanism is that hypoxia promoted by impaired vessel perfusion not only induces the secretion of cytokines such as TGF◻, VEGF, IL-10 to increase the recruitment of immunosuppressive cells but also upregulates the expression of CTLA-4 or LAG3 on regulatory T cells and PD-L1 on myeloid-derived suppressor cells, tumor-associated macrophages and tumor cells^15,38,39^. Treatment with dual anti-CTLA-4 and anti-PD1 therapy induced tumor vessel normalization through a dynamic process that involves various immune populations at different stages of tumor development^15,17^. Although the precise mechanism remains unknown, it is reported that type 1 helper T cells play a crucial role in vascular normalization by immune checkpoint blockade^17^. An increase in tumor vascularity during immune rejection was detected by DCE-MRI in a E.G7-OVA tumor-bearing mice model which undergoes spontaneous regression^40^. As for ICB therapy, there was one report showing that DCE-MRI revealed no alteration in relative blood distribution volume in anti-PD-L1 and anti-CTLA-4 AB treated CT26 tumor^41^. In the current study, K^trans^ increased in treated MC38 and B16.F10 tumors relative to control tumors and the increase was parallel to the treatment effect. Thus, K^trans^ can potentially reflect early treatment responses. IHC of CD31 suggested increased vascular vessels in ICB treated tumor. However, it should be mentioned that increased vascular vessels does not simply mean vascular normalization because majority of the reports especially related to anti-anigiogenesis agent show decreased CD31 staining by this therapy. In this respect, one previous report gives evidence of vascular normalization by ICB therapy as they showed that functional vessels covered by pericyte increased by dual blockade of PD-1 and CTLA-4^17^, while those decreased in above-referenced another report^41^. Thus, it is still controversial and further investigation is needed for DCE-MRI to be used as imaging method to detect ICB therapy.

In conclusion, we performed multi-modal imaging to detect the early response to immune checkpoint blockade therapy. DNP ^13^C MRI with [1,4-13C2] fumarate showed enhanced production of malate reflecting necrotic tumor cell death after the therapy. DCE-MRI showed increased intratumor permeability in treated tumor. The magnitude of both measures paralled the effectiveness of ICB treatment considered on a per model basis. Collectively, these findings suggest that these imaging methods can provide predictive biomarkers for effect of immune checkpoint blockade and better understanding metabolic and physiologic profile of cancer.

## Methods

### Mice and tumor

The animal experiments were conducted according to a protocol approved by the Animal Research Advisory Committee of the NIH (RBB-159-3E) in accordance with the National Institutes of Health Guidelines for Animal Research. Female C57BL/6 (B6) mice were supplied by the Frederick Cancer Research Center, Animal Production (Frederick, MD) and housed in a specific pathogen-free environment and used at an age of 8-12 week. MC38 colon adenocarcinoma and B16.F10 melanoma were used as sensitive or less sensitive models to immune checkpoint blockade therapy. MC38 was purchased from Kerafast (Boston, MA). B16.F10 was used from the frozen stock in our lab and tested in Feb 2020 and authentificated by IDEXX RADIL (Columbia, MO) using a panel of microsatellite markers. Molecular testing of cell lines for multiple pathogens, including mycoplasma, was performed at the time of receipt and prior to in vivo studies. Both cell lines were maintained in DMEM containing 4.5 g/L glucose supplemented with 10% fetal calf serum and antibiotics.

### In vivo checkpoint blockade treatment

For the in vivo treatment model, 1 × 10^5^ MC38 tumor cells or 2 × 10^5^ B16.F10 tumor cells were inoculated s.c. into the right leg of B6 mice. Tumor bearing mice were injected i.p. with 200 μg of ◻PD-L1 (10F.9G2) or αPD-1 (RMP1-14) or αCTLA-4 (9H10) antibody or with isotype control antibodies (all from BioXcell) on days 9, 12, and 15 post tumor inoculation at the size of approximately 100-150 mm^3^. Imaging experiments were performed after 1^st^ to 3^rd^ injection of antibodies.

### In vitro treatment

For in vitro treatment assay, tumor cells were cultured with 100 U/ml of recombinant murine IFN-◻ for 48h followed by 10 μg/ml ◻PD-L1 antibody treatment for an additional 24h before assay.

### Flow cytometry

To generate activated tumor-draining lymph node (TDLN) cells, B6 mice were inoculated s.d. with 1 × 10^5^ MC38 tumor cells on both flanks to stimulate TDLNs. 13 days later, TDLNs (inguinal) were harvested, and single-cell suspensions were prepared mechanically. To analyze tumor infiltrating lymphocytes, single-cell suspensions were prepared from solid tumors by digestion with a mixture of 0.1 % collagenase, 0.01 % DNase, and 2.5 U/ml hyaluronidase for 3 h at 37 deg C. FITC-conjugated mAbs against CD4 (RM4-5); PE-conjugated mAbs against CD3; Cy-chrome-conjugated mAbs against CD8 (53-6.7); isotype-matched mAbs were purchased from BD Biosciences. The cell surface phenotypes were determined by direct immunofluorescence staining with conjugated mAbs and analyzed using FACS Calibur (BD Biosciences, San Jose, CA). Digested cells were stained with αCD3, αCD8, and αCD4 antibody. Data were collected on FACS Calibur flow cytometer. Tumor infiltrating lymphocytes were identified and gated on forward scatter vs side scatter plot.

### Western blot

In vitro treated tumor cells on the dish were lysed in radioimmunoprecipitation assay buffer (RIPA buffer, Thermo Scientific) supplemented with protease and phosphatase inhibitors (Roche). Protein concentrations were measured by bicinchoninic acid assay (BCA protein assay, Thermo Fisher Scientific). Glut-1, Hexokinase-2, lactate dehydrogenase A (LDHA), monocarboxylate transporter 1 (MCT1), MCT4 proteins were separated on 4 % to 20 % Tris-Glycine gel (Life Technologies) by SDS-PAGE and were transferred to nitrocellulose membrane. The membranes were blocked for 1 h in blocking buffer (3 % nonfat dry milk in 0.1 % Tween 20/TBS), which was then replaced by the primary antibody (1:500-1:1000), diluted in blocking buffer, and then incubated for 1 h at room temperature. The membranes were then washed three times in washing buffer (0.1 % Tween 20/TBS). The primary antibody was detected using the appropriate horseradish peroxidase conjugated secondary antibody and measured by the Fluor Chem HD2 chemiluminescent imaging system (Alpha Innotech Corp.). Density values for each protein were normalized to actin or HSC70.

### Seahorse metabolism assay

Extracellular acidification rate (ECAR) was analyzed on a XF96 Extracellular Flux Analyzer using XF Glycolysis Stress Test Kit (Agilent Technology, Santa Clara, CA) in accordance with the manufacturer’s instructions. Briefly, first, in vitro treated cells plated on 96 well plate were incubated in XF base medium without glucose or pyruvate. Glucose was injected and the ECAR is measured as the rate of glycolysis under basal conditions. Then, oligomycin, an ATP synthase inhibitor was injected and ECAR was monitored (maximum glycolytic capacity). Finally, 2-deoxy-glucose (2-DG), a glucose analog, that inhibits glycolysis through competitive binding to glucose hexokinase, was injected to confirm that the ECAR produced in the experiment was due to glycolysis.

### Histological assessment

Tumor tissues were excised, frozen with Tissue-Tek O.C.T. compound (Sakura Finetek USA Inc.) by ultra-cold ethanol and sectioned (10 mm) using a cryostat, with the sections being thaw-mounted on glass slides. After fixing with 4 % paraformaldehyde, sections were treated with cold acetone for 15 minutes. After blocking nonspecific-binding sites on sections with Protein Block Serum-Free reagent (Dako North America Inc.) for 30 minutes, the slides were covered by CD3 antibody (BD Biosciences; 1:250) combined with CD8 antibody (Abcam, Inc.; 1:250) overnight at 4 C. The sections were then incubated with Alexa Fluor 488 antimouse and the Alexa Fluor 546 F(ab’)2 fragment of goat antirabbit IgG (HþL) (Invitrogen; 1:2,000) for 1 h at room temperature, before being mounted with Prolong Gold antifade reagent with DAPI (Invitrogen). CD31 was stained with same procedure using CD31 antibody (BD Biosciences; 1:250) and Alexa Fluor 488 antirat (Invitrogen; 1:250). The sections were also stained with hematoxylin and eosin and mounted on permount for histological observation. The stained slides were scanned using a BZ-9000 microscope (Keyence), and the immunostain-positive area was quantified using ImageJ software (downloaded from https://imagej.nih.gov/ij/).

### Hyperpolarized ^13^C MRI

Details of the hyperpolarization procedure were reported previously^22,23^. Briefly, [1-^13^C] pyruvic acid (30 μL) or [1,4-^13^C_2_] fumaric acid (2.5 M in 30 μL deuterated DMSO), containing 15 mmol/L Ox063 and 2.5 mmol/L of the gadolinium chelate ProHance (BraccoDiagnostics) was polarized in Hypersense DNP polarizer (Oxford Instruments). After the polarization reached 80 % of the plateau value, the hyperpolarized sample was rapidly dissolved in 4.5 mL of a superheated alkaline buffer consisted of 40 mmol/L HEPES, NaOH, and 100 mg/L EDTA. Hyperpolarized [1-^13^C] pyruvate or [1,4-^13^C_2_] fumarate solution was rapidly injected intravenously through a catheter placed in the tail vein of each mouse (12 mL/g body weight). Hyperpolarized ^13^C MRI studies were performed on a 3 T dedicated MR Solutions animal scanner (MR SOLUTIONS Ltd., Boston, MA) using a 17 mm home-built ^13^C solenoid coil placed inside of a saddle coil tuned to ^1^H frequency. Both ^1^H and ^13^C were tuned and matched and anatomical image was taken after shimming on proton. ^13^C spectra was acquired every 1 s for 240 s from the whole leg including each tumor. The repetition time, spectral width, flip angle, and number of average were 1000 ms, 3300 Hz, 10°, and 1, respectively.

### Fumarase activity assay

A serum sample was obtained from isotype control antibody treated mice and from αPD-L1 and αCTLA-4 antibodies treated mice one day after 2^nd^ injection. Fumarase activity was measured with Fumarase Assay Kit (BioAssay Systems, Hayward, CA) in accordance with the manufacturer’s instruction manual.

### DCE-MRI of Gd-DTPA

DCE-MRI studies were performed on a 1 T scanner (Bruker BioSpin MRI GmbH). T1-weighted fast low-angle shot (FLASH) images were obtained with TR = 156 ms; TE = 4 ms; flip angle = 45°; four slices; 0.44 × 0.44 mm resolution; 15-second acquisition time per image; and 45 repetitions. Gd-DTPA solution (4 mL/g of body weight of 50 mmol/L Gd-DTPA) was injected through a tail vein cannula 1 minutes after the start of the dynamic FLASH sequence. To determine the local concentrations of Gd-DTPA, T1 maps were calculated from three sets of Rapid Imaging with Refocused Echoes (RARE) images obtained with TR = 300, 600, 1000, and 2,000 ms, with the acquisitions being made before running the FLASH sequence.

### Statistical analysis

The significance of the differences between groups was analyzed using the Wilcoxon rank sum test or the Student t test. Kaplan-Meier curves were constructed for the survival of mice; the differences between groups were identified using the log-rank test. A two-tailed *p* value, 0.05 was considered significant. All experiments were repeated at least twice.

## Acknowledgements

This research was supported by the Intramural Research Program, Center for Cancer Research, NCI, NIH (grant number Z01BC010477). Y.S. is a JSPS (Japan Society for the Promotion of Science) -NIH Research Fellow.

## Author Contributions

Y.S. and S.K. conceptualized, designed, and performed the study. Y.S., S.K., and J.R.B. performed data analysis. Y.S., S.K., J.R.B., K.Y., and M.C.K. interpreted the results. Y.S. and S.K. wrote the original manuscript. J.R.B., J.B.M. and M.C.K. reviewed and edited the manuscript. M.C.K. supervised the project.

## References

1 Mellman, I., Coukos, G. & Dranoff, G. Cancer immunotherapy comes of age. Nature 480, 480–489, doi:10.1038/nature10673 (2011).

2 Wei, S. C., Duffy, C. R. & Allison, J. P. Fundamental Mechanisms of Immune Checkpoint Blockade Therapy. Cancer discovery 8, 1069–1086, doi:10.1158/2159-8290.Cd-18-0367 (2018).

3 Sharma, P. & Allison, J. P. Immune checkpoint targeting in cancer therapy: toward combination strategies with curative potential. Cell 161, 205–214, doi:10.1016/j.cell.2015.03.030 (2015).

4 Nishino, M., Ramaiya, N. H., Hatabu, H. & Hodi, F. S. Monitoring immune-checkpoint blockade: response evaluation and biomarker development. Nature reviews. Clinical oncology 14, 655–668, doi:10.1038/nrclinonc.2017.88 (2017).

5 Kwak, J. J., Tirumani, S. H., Van den Abbeele, A. D., Koo, P. J. & Jacene, H. A. Cancer immunotherapy: imaging assessment of novel treatment response patterns and immune-related adverse events. Radiographics : a review publication of the Radiological Society of North America, Inc 35, 424–437, doi:10.1148/rg.352140121 (2015).

6 Wolchok, J. D. et al. Guidelines for the evaluation of immune therapy activity in solid tumors: immune-related response criteria. Clinical cancer research : an official journal of the American Association for Cancer Research 15, 7412–7420, doi:10.1158/1078-0432.Ccr-09-1624 (2009).

7 Weiss, J., Notohamiprodjo, M., Bedke, J., Nikolaou, K. & Kaufmann, S. Imaging response assessment of immunotherapy in patients with renal cell and urothelial carcinoma. Current opinion in urology 28, 35–41, doi:10.1097/mou.0000000000000463 (2018).

8 Butterfield, L. H. et al. Recommendations from the iSBTc-SITC/FDA/NCI Workshop on Immunotherapy Biomarkers. Clinical cancer research : an official journal of the American Association for Cancer Research 17, 3064–3076, doi:10.1158/1078-0432.Ccr-10-2234 (2011).

9 Galon, J. & Bruni, D. Approaches to treat immune hot, altered and cold tumours with combination immunotherapies. Nature reviews. Drug discovery 18, 197–218, doi:10.1038/s41573-018-0007-y (2019).

10 MacIver, N. J., Michalek, R. D. & Rathmell, J. C. Metabolic regulation of T lymphocytes. Annual review of immunology 31, 259–283, doi:10.1146/annurev-immunol-032712-095956 (2013).

11 Sukumar, M., Roychoudhuri, R. & Restifo, N. P. Nutrient Competition: A New Axis of Tumor Immunosuppression. Cell 162, 1206–1208, doi:10.1016/j.cell.2015.08.064 (2015).

12 Chang, C. H. et al. Metabolic Competition in the Tumor Microenvironment Is a Driver of Cancer Progression. Cell 162, 1229–1241, doi:10.1016/j.cell.2015.08.016 (2015).

13 Hope, H. C. & Salmond, R. J. Targeting the tumor microenvironment and T cell metabolism for effective cancer immunotherapy. European journal of immunology 49, 1147–1152, doi:10.1002/eji.201848058 (2019).

14 Brand, A. et al. LDHA-Associated Lactic Acid Production Blunts Tumor Immunosurveillance by T and NK Cells. Cell metabolism 24, 657–671, doi:10.1016/j.cmet.2016.08.011 (2016).

15 Huang, Y. et al. Improving immune-vascular crosstalk for cancer immunotherapy. Nature reviews. Immunology 18, 195–203, doi:10.1038/nri.2017.145 (2018).

16 Huang, Y., Stylianopoulos, T., Duda, D. G., Fukumura, D. & Jain, R. K. Benefits of vascular normalization are dose and time dependent--letter. Cancer research 73, 7144–7146, doi:10.1158/0008-5472.Can-13-1989 (2013).

17 Tian, L. et al. Mutual regulation of tumour vessel normalization and immunostimulatory reprogramming. Nature 544, 250–254, doi:10.1038/nature21724 (2017).

18 Kaira, K. et al. Metabolic activity by (18)F-FDG-PET/CT is predictive of early response after nivolumab in previously treated NSCLC. European journal of nuclear medicine and molecular imaging 45, 56–66, doi:10.1007/s00259-017-3806-1 (2018).

19 Cho, S. Y. et al. Prediction of Response to Immune Checkpoint Inhibitor Therapy Using Early-Time-Point (18)F-FDG PET/CT Imaging in Patients with Advanced Melanoma. Journal of nuclear medicine : official publication, Society of Nuclear Medicine 58, 1421–1428, doi:10.2967/jnumed.116.188839 (2017).

20 Bier, G. et al. CT imaging of bone and bone marrow infiltration in malignant melanoma--Challenges and limitations for clinical staging in comparison to 18FDG-PET/CT. European journal of radiology 85, 732–738, doi:10.1016/j.ejrad.2016.01.012 (2016).

21 Kong, B. Y. et al. Residual FDG-PET metabolic activity in metastatic melanoma patients with prolonged response to anti-PD-1 therapy. Pigment cell & melanoma research 29, 572–577, doi:10.1111/pcmr.12503 (2016).

22 Matsumoto, S. et al. In vivo imaging of tumor physiological, metabolic, and redox changes in response to the anti-angiogenic agent sunitinib: longitudinal assessment to identify transient vascular renormalization. Antioxidants & redox signaling 21, 1145–1155, doi:10.1089/ars.2013.5725 (2014).

23 Matsumoto, S. et al. Metabolic and Physiologic Imaging Biomarkers of the Tumor Microenvironment Predict Treatment Outcome with Radiation or a Hypoxia-Activated Prodrug in Mice. Cancer research 78, 3783–3792, doi:10.1158/0008-5472.Can-18-0491 (2018).

24 Gallagher, F. A. et al. Production of hyperpolarized [1,4-13C2]malate from [1,4-13C2]fumarate is a marker of cell necrosis and treatment response in tumors. Proceedings of the National Academy of Sciences of the United States of America 106, 19801–19806, doi:10.1073/pnas.0911447106 (2009).

25 Kishimoto, S. et al. Molecular imaging of tumor photoimmunotherapy: Evidence of photosensitized tumor necrosis and hemodynamic changes. Free radical biology & medicine 116, 1–10, doi:10.1016/j.freeradbiomed.2017.12.034 (2018).

26 Postow, M. A. et al. Nivolumab and ipilimumab versus ipilimumab in untreated melanoma. The New England journal of medicine 372, 2006–2017, doi:10.1056/NEJMoa1414428 (2015).

27 Larkin, J. et al. Combined Nivolumab and Ipilimumab or Monotherapy in Untreated Melanoma. The New England journal of medicine 373, 23–34, doi:10.1056/NEJMoa1504030 (2015).

28 Hodi, F. S. et al. Nivolumab plus ipilimumab or nivolumab alone versus ipilimumab alone in advanced melanoma (CheckMate 067): 4-year outcomes of a multicentre, randomised, phase 3 trial. The Lancet. Oncology 19, 1480–1492, doi:10.1016/s1470-2045(18)30700-9 (2018).

29 Juneja, V. R. et al. PD-L1 on tumor cells is sufficient for immune evasion in immunogenic tumors and inhibits CD8 T cell cytotoxicity. The Journal of experimental medicine 214, 895–904, doi:10.1084/jem.20160801 (2017).

30 Patsoukis, N. et al. PD-1 alters T-cell metabolic reprogramming by inhibiting glycolysis and promoting lipolysis and fatty acid oxidation. Nature communications 6, 6692, doi:10.1038/ncomms7692 (2015).

31 Allison, K. E., Coomber, B. L. & Bridle, B. W. Metabolic reprogramming in the tumour microenvironment: a hallmark shared by cancer cells and T lymphocytes. Immunology 152, 175–184, doi:10.1111/imm.12777 (2017).

32 Siska, P. J. & Rathmell, J. C. T cell metabolic fitness in antitumor immunity. Trends in immunology 36, 257–264, doi:10.1016/j.it.2015.02.007 (2015).

33 Bohn, T. et al. Tumor immunoevasion via acidosis-dependent induction of regulatory tumor-associated macrophages. Nature immunology 19, 1319–1329, doi:10.1038/s41590-018-0226-8 (2018).

34 Bohndiek, S. E., Kettunen, M. I., Hu, D. E. & Brindle, K. M. Hyperpolarized (13)C spectroscopy detects early changes in tumor vasculature and metabolism after VEGF neutralization. Cancer research 72, 854–864, doi:10.1158/0008-5472.Can-11-2795 (2012).

35 Mignion, L. et al. Monitoring chemotherapeutic response by hyperpolarized 13C-fumarate MRS and diffusion MRI. Cancer research 74, 686–694, doi:10.1158/0008-5472.Can-13-1914 (2014).

36 Martinez-Lostao, L., Anel, A. & Pardo, J. How Do Cytotoxic Lymphocytes Kill Cancer Cells? Clinical cancer research : an official journal of the American Association for Cancer Research 21, 5047–5056, doi:10.1158/1078-0432.Ccr-15-0685 (2015).

37 Vandenabeele, P., Galluzzi, L., Vanden Berghe, T. & Kroemer, G. Molecular mechanisms of necroptosis: an ordered cellular explosion. Nature reviews. Molecular cell biology 11, 700–714, doi:10.1038/nrm2970 (2010).

38 Barsoum, I. B., Smallwood, C. A., Siemens, D. R. & Graham, C. H. A mechanism of hypoxia-mediated escape from adaptive immunity in cancer cells. Cancer research 74, 665–674, doi:10.1158/0008-5472.Can-13-0992 (2014).

39 Doedens, A. L. et al. Hypoxia-inducible factors enhance the effector responses of CD8(+) T cells to persistent antigen. Nature immunology 14, 1173–1182, doi:10.1038/ni.2714 (2013).

40 Hu, D. E., Beauregard, D. A., Bearchell, M. C., Thomsen, L. L. & Brindle, K. M. Early detection of tumour immune-rejection using magnetic resonance imaging. British journal of cancer 88, 1135–1142, doi:10.1038/sj.bjc.6600814 (2003).

41 Fiegle, E. et al. Dual CTLA-4 and PD-L1 Blockade Inhibits Tumor Growth and Liver Metastasis in a Highly Aggressive Orthotopic Mouse Model of Colon Cancer. Neoplasia (New York, N.Y.) 21, 932–944, doi:10.1016/j.neo.2019.07.006 (2019).

