## Supplementary figures and images for "Detecting Early Response to Immune Checkpoint Blockade by Multimodal Molecular Imaging"

### SI Figure 1

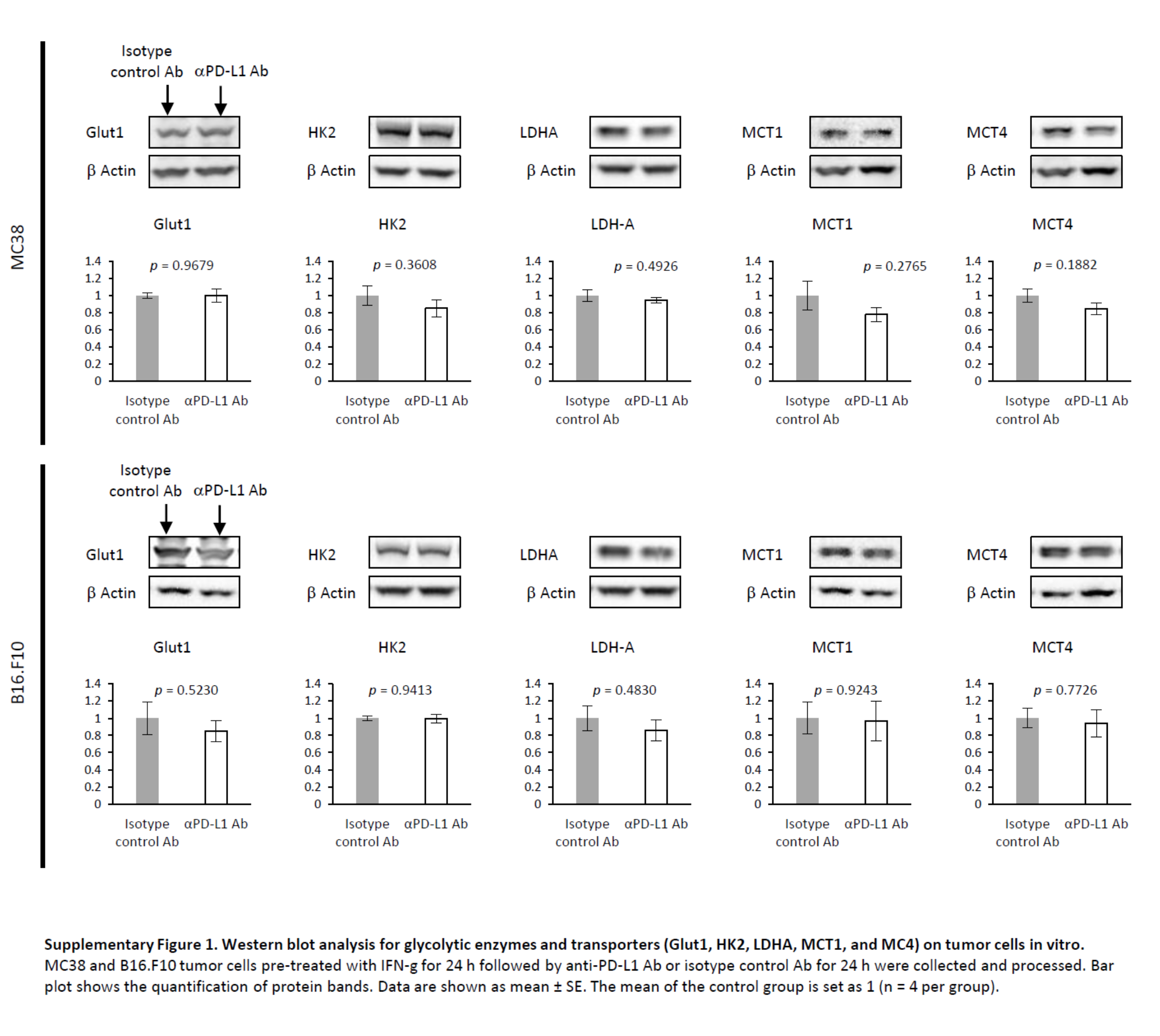
